# Endosomal hyper-acidification via proton-activated chloride channel deletion in neurons impairs AMPA receptor endocytosis and LTD

**DOI:** 10.1101/2024.09.27.615471

**Authors:** Kevin H. Chen, Junhua Yang, Bian Liu, Chaohua Jiang, Nicholas Koylass, Zhe Zhang, Shuying Sun, Richard Huganir, Zhaozhu Qiu

**Author notes:** Corresponding author. Zhaozhu Qiu. **Author Contributions:** K.H.C. performed the majority of experiments. K.H.C. and Z.Q. designed the study and wrote the manuscript with input from all authors. J.Y., C.J., and N.K. performed electrophysiological recordings. B.L. performed imaging experiments. Z.Z., S.S. and R.H. provided critical reagents. **Competing Interest Statement:** The authors declare that they have no competing interests. **Classification:** Biological Sciences, Neuroscience.

## Abstract

Endosomal homeostasis is critical for neuronal function, including the post-synaptic trafficking of α-amino-3-hydroxy-5-methyl-4-isoxazolepropionic acid receptors (AMPARs). Dynamic AMPAR trafficking is a major component of synaptic plasticity, such as hippocampal long-term potentiation (LTP) and long-term depression (LTD) and is thought to be required for learning and memory. Dramatic alteration of endosomal pH has been reported to negatively affect synaptic transmission and neural development, but the underlying mechanisms by which pH is involved in AMPAR trafficking are unclear. Here, we show that the proton-activated chloride (PAC) channel localizes to early and recycling endosomes along neuronal dendrites and prevents hyper-acidification of endosomes. To directly measure AMPAR endocytosis, we used a new method to assess LTD using HaloTag-GluA2 and found that the loss of PAC reduces AMPAR internalization during chemical LTD in primary neurons, while AMPAR trafficking in unstimulated cells or during chemical LTP is unaffected. Consistently, pyramidal neuron-specific PAC knockout mice had impaired hippocampal LTD, but not LTP, and performed poorly in the Morris water maze reversal test, exhibiting an inability to adapt to changing environments (also referred to as behavioral flexibility). We conclude that proper maintenance of pH by PAC is important during LTD to regulate AMPAR trafficking in a manner critical for animal physiology and behavior.

**Summary:** The ability to adapt to changing environments stems from the plasticity of neurons, which can modulate their synaptic strength in response to neural activity. We discovered a novel mechanism by which an endosomal proton-activated chloride channel (PAC) is involved in synaptic weakening, or long-term depression (LTD). To improve tools used to study LTD, we developed a live-cell imaging assay to directly observe α-amino-3-hydroxy-5-methyl-4-isoxazolepropionic acid receptor (AMPAR) endocytosis. PAC-deficient neurons have hyper-acidified endosomes and fail to endocytose AMPARs during LTD. Neuron-specific PAC knockout mice have impaired hippocampal LTD and fail to adapt to changes in their environment. The role of endosomal pH in synaptic function is understudied, and our results provide a novel mechanism whereby PAC can affect synaptic LTD.

## Introduction

Synaptic plasticity, the ability of individual synapses to modulate their connective strength based on patterns of neural activity, is thought to be the molecular basis of neural plasticity and underlies complex behaviors such as learning and memory ^1^. Long-term potentiation (LTP) and long-term depression (LTD) are well-established models of long-lasting synaptic plasticity, and at glutamatergic synapses, involve the modulation of post-synaptic α-amino-3-hydroxy-5-methyl-4-isoxazolepropionic acid receptors (AMPARs) either through dynamic AMPAR trafficking at the post-synaptic membrane or modulation of channel properties and binding proteins ^2^. AMPARs can be trafficked through synapses, extrasynaptic membranes, and the endocytic pathway, allowing neurons to regulate the connectivity and strength of glutamatergic synapses. LTP occurs by enhancing the delivery of AMPARs to the post-synaptic membrane from extrasynaptic and endosomal membranes ^3–5^. Conversely, LTD reduces AMPAR content at the post-synaptic density (PSD) through diffusion and endocytosis, followed by either sequestration in recycling endosomes or degradation through the endolysosomal pathway ^6^. While many mechanisms are involved in synaptic plasticity, the expression of excitatory synaptic plasticity through AMPARs is inextricably linked to the endosomal pathway ^6–9^.

Disrupted endosomal function, which could lead to endosomal alkalinization or hyper-acidification, can impair synaptic plasticity. Indeed, direct endosomal alkalinization by an optogenetically-activated proton leak channel abolishes LTD in the cerebellum ^10^. Consistent with these findings, aberrations in AMPAR trafficking and synaptic plasticity have been shown to occur during genetic ablation of various endosomal adaptor proteins that interact with AMPARs, such as protein interacting with C kinase 1 (PICK1) ^11–13^, synaptotagmin-3 ^14^, GRIP-associated protein 1 (GRASP1) ^15^, or the Rab family of small GTPases, which compartmentalize endosomal membranes and regulate endosome identity ^16–18^. Moreover, dysfunction of multiple ion transporters, including the V-type ATPase, chloride transporters (CLCs), and endosomal sodium-proton exchangers (NHEs), which regulation of ion homeostasis and pH in the endosomal lumen, are associated with neuropsychological impairments, including Alzheimer’s disease, intellectual disability, and autism spectrum disorders ^19–22^. Taken together, proper synaptic function depends on the trafficking and signaling roles of the endolysosomal system. However, little is known how endosome hyper-acidification directly affects AMPAR trafficking during synaptic plasticity.

Through an unbiased screen, we have recently revealed the molecular identity of the proton-activated chloride (PAC) channel ^23^. PAC is ubiquitously expressed and primarily localized to endosomes in various cell lines. Endosomal PAC becomes activated in the low pH environment of endosomes and functions as a chloride release channel to lower luminal [Cl^-^] and prevent hyper-acidification ^24^. In macrophages, PAC also regulates macropinosome volume resolution ^25^. Under pathological acidosis conditions, the plasma membrane PAC channel has been shown to mediate acid-induced cytotoxicity and increase susceptibility to ischemic stroke in mice ^23,26^. Despite much progress in studying the biophysical and cellular roles of PAC, its function has not been established in the context of AMPAR trafficking and synaptic plasticity in the brain.

Here, we developed a chemogenetic approach to longitudinally visualize AMPAR endocytosis and determined that loss of functional PAC channels in neurons impairs endosomal pH homeostasis and disrupts the AMPAR internalization pathway required for hippocampal LTD. Moreover, we found that PAC impairment causes changes in animal behavior during tasks requiring behavioral flexibility, which is the ability to adapt to changing environments. PAC deletion specifically altered activity-dependent AMPAR endocytosis without affecting LTP or basal AMPAR trafficking, revealing a distinct role for PAC-dependent endosomal acidification in synaptic plasticity.

## Results

### PAC Localizes to Early and Recycling Endosomes, but not Synaptic Vesicles

PAC localization in neurons has not been established. To explore the subcellular location of PAC, we transfected primary hippocampal rat neurons with a human isoform of PAC (hPAC) and stained neurons using a validated antibody developed previously ^24^. Neurons co-expressed cytosolic mVenus and were immunolabeled with anti-hPAC and MAP2 antibodies. Axons and dendrites were distinguished though cellular morphology and fluorescence signals from mVenus and MAP2, a dendrite marker, where axons are MAP2-negative/mVenus-positive and dendrites are MAP2-positive. Using this approach, we observed PAC puncta in the cell soma, axons, and dendrites (Fig. S1A).

In dendrites, PAC mainly colocalizes with the early endosome marker EEA1 and the recycling endosome marker syntaxin13 (Fig. 1A, B). PAC does not localize to the PSD marked by Homer1 (Fig. 1A, B), but is positioned adjacent to the PSD, reminiscent of endocytic zones ^5,9,27^. To label presynaptic proteins, we used lentiviral transduction of hPAC with a high transduction efficiency to express PAC in nearly all neurons but did not find PAC colocalization with pre-synaptic synapsin1 or with Vglut1-containing synaptic vesicles (Fig. 1A, B). These results indicate that PAC localizes mainly to early and recycling endosomes in dendrites.

**Figure 1.**
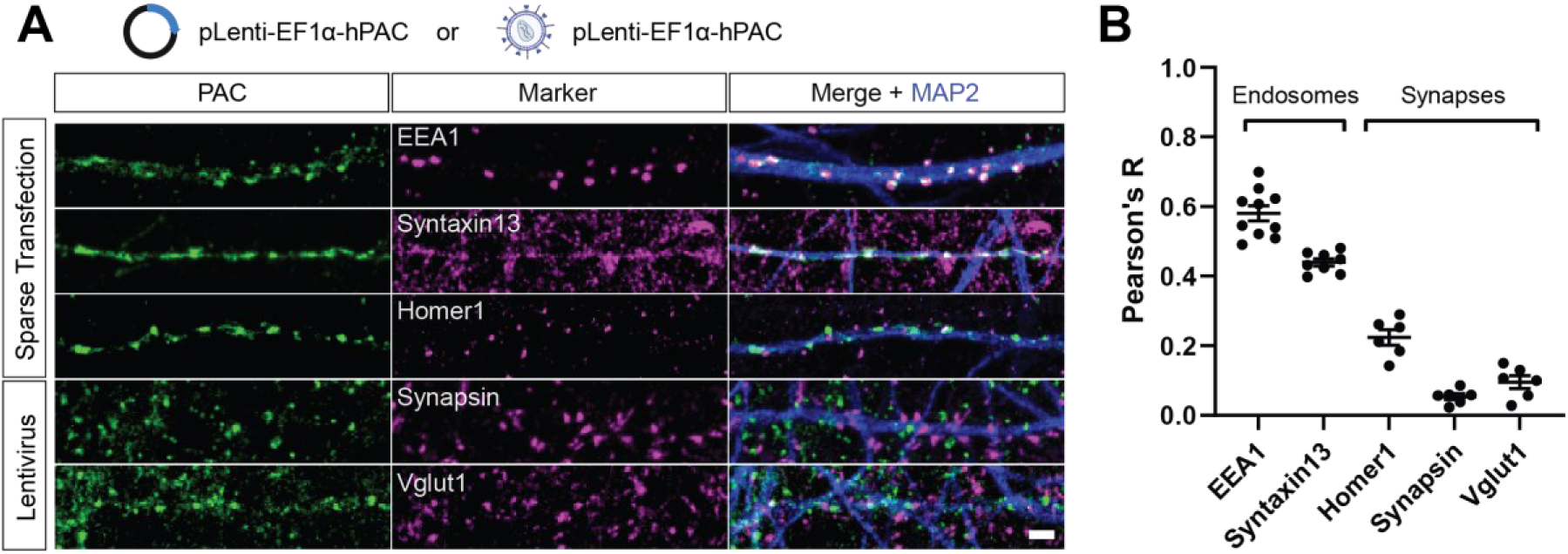
PAC localizes to dendritic endosomes in neurons. **(A)** Co-immunofluorescence labeling of PAC and cellular markers in rat primary hippocampal neurons. Sparse transfection was used to visualize post-synaptic neurons, and lentiviral transduction was used to visualize pre-synaptic markers. Scale bar 2 µm. **(B)** Quantification (mean ± SEM) of colocalization using Pearson’s R for PAC and various markers. PAC localization is correlated with EEA1, Syntaxin13, and is not closely correlated with Homer1, Synapsin, and Vglut1. Points represent 1 neuron.

In addition to rat primary hippocampal neurons, we also investigated PAC localization in mouse primary hippocampal neurons and human I3 IPSC-derived neurons (IPSNs), the latter of which allowed us to examine the endogenous expression of PAC in human cells (Fig. S1). Consistent with our data from rat neurons, PAC in mouse hippocampal neurons colocalizes with EEA1 and syntaxin13 but does not localize to the post-synaptic density protein PSD-95 or lysosomal LAMP2 (Fig. S1B). In human IPSNs, we detected endogenous PAC expression and colocalization with EEA1 (Fig. S1C, D). We validated the staining specificity by knocking down endogenous PAC with CRISPRi and sgRNAs targeting its transcriptional start site, which abolished PAC signal in PAC knockdown (KD) neurons (Fig. S1C, D). In agreement with previous reports of PAC localization, our results demonstrate that PAC localizes to early and recycling endosomes but not to synapses, synaptic vesicles, or lysosomes.

### PAC regulates endosomal pH in neurons

We assessed the role of PAC in primary hippocampal and cortical neurons by expressing either control shRNA or PAC-specific shRNA to generate control and PAC KD neurons, respectively ^28^. To further validate our PAC-specific shRNA, we developed a PAC antibody targeting the rodent PAC isoform and used Western blotting to measure total PAC levels (Fig. S2). Consistently, PAC protein was markedly reduced in PAC KD neurons compared to control neurons. PAC regulates endosomal pH by controlling luminal Cl^-^ levels in several cancer cell lines, but this has not been explored in neurons. To measure endosomal pH in primary neurons, we used ratiometric pHluorin (RpH), a GFP mutant with two pH-dependent excitation peaks (405 nm/488 nm), where the ratio of these peaks is proportional to ambient pH ^29^. To target RpH to endosomes, we fused the protein to the extracellular domain of the transferrin receptor (TfR) to generate a TfR-RpH fusion protein, which co-localizes with PAC in endosomes (Fig. 2A, B). TfR-RpH fluorescence ratios of endosomes in hippocampal neurons co-expressing control or PAC shRNAs was measured first under physiological conditions in artificial cerebrospinal fluid (ACSF). Then, neurons were treated with pH calibration solutions containing the ionophore monensin, which equilibrates H^+^ ions across cellular membranes (Fig. 2C). As expected, the endosomal 405/488 nm excitation ratio decreased as pH decreased from 7.5 to 5.5. Fluorescence intensities of endosomes under the pH calibration solutions were plotted to generate a standard curve, which was used to calculate the physiological pH of endosomes measured in ACSF. We observed a significant decrease in endosomal pH in PAC KD (pH = 5.997 ± 0.030) cells compared to control (pH = 6.493 ± 0.0785) (Fig. 2D), indicating that PAC prevents hyper-acidification in neuronal endosomes.

**Figure 2.**
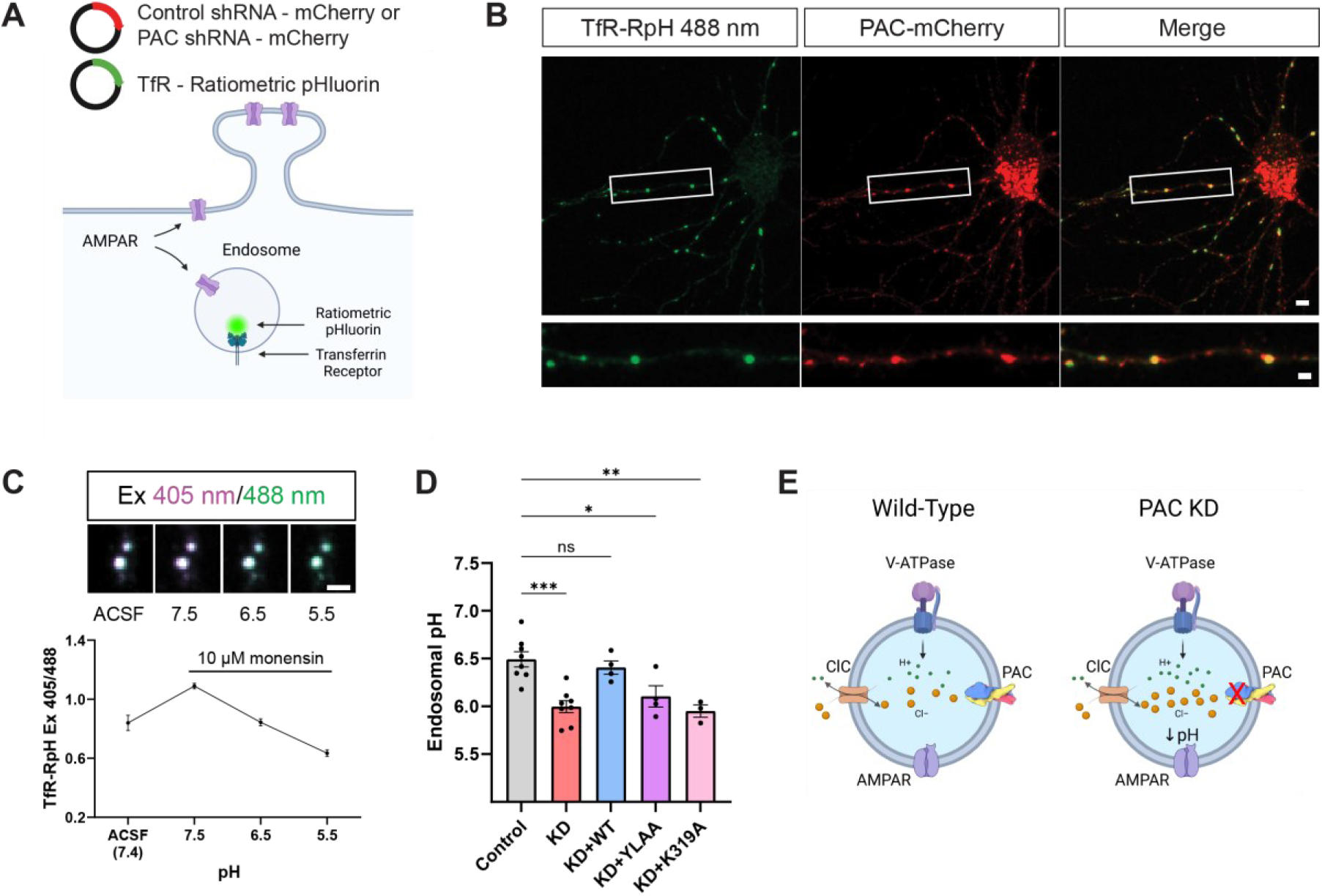
PAC regulates endosomal pH in dendrites. **(A)** Schematic of the ratiometric endosomal pH reporter TfR-RpH (transferrin receptor – ratiometric pHluorin) and co-transfection with control or PAC shRNA. **(B)** Representative live-neuron expressing TfR-RpH pH reporter and colocalization with PAC. Scale bars: upper, 10 µm; lower, 2 µm. **(C)** Fluorescence intensity measurements of individual endosomes under basal conditions and in pH 7.5 – 5.5 calibration buffers. Scale bar 2 µm. **(D)** Ratiometric measurement of endosomal pH in neurons treated with control shRNA (ctrl, 6.475 ± 0.063, n = 58), PAC shRNA (KD, 6.021 ± 0.044, n = 68), or KD plus PAC WT (KD+WT, 6.391 ± 0.073, n = 50), PAC YLAA mutant (KD+YLAA, 6.050 ± 0.091, n = 47), or PAC K319A mutant (KD+K319A, 6.049 ± 0.088, n = 47). Points represent 1 experiment and n represents neurons. Error bars represent mean ± SEM. Two-way ANOVA with Tukey correction for multiple comparisons. ***p < 0.001, **p < 0.01, *p < 0.05, ns = not significant. **(E)** Model of how PAC regulates endosomal pH through pH-dependent chloride efflux.

To determine which functional domains of the PAC channel are necessary for regulating endosomal pH, we employed a molecular replacement strategy where hippocampal neurons were transfected with an shRNA targeting PAC plus shRNA-resistant wild-type (WT) hPAC, endocytosis defective mutant Y10A/L13A (YLAA), or channel dead mutant K319A (Fig. 2D). WT and K319A PAC colocalized with endosomes, whereas the YLAA mutant was broadly distributed at the plasma membrane (Fig. S3A). Whole-cell patch clamp recordings from K319A-expressing neurons revealed a large reduction in proton-activated chloride channel activity (Fig. S3B). Overexpression of WT hPAC in PAC shRNA-transfected neurons increased endosomal pH (pH = 6.406 ± 0.070), thereby rescuing the hyper-acidification phenotype. However, neither the YLAA (pH = 6.104 ± 0.112) nor the K319A (pH = 5.951 ± 0.065) mutants rescued endosomal pH in PAC KD neurons. These results demonstrate that PAC prevents endosomal hyper-acidification, and that endosomal pH regulation depends on the proper localization and activity of the channel.

### PAC does not affect AMPAR trafficking in unstimulated neurons or during chemical LTP

The phenotype in PAC KD neurons provides a rare opportunity to examine the potential effects of endosomal hyper-acidification on neuronal function, which has not been explored previously. PAC KD did not disrupt the expression levels of many membrane and endosomal proteins in primary neurons (Fig. S2). Protein levels of Na/K-ATPase, GluA1, GluA2, PSD-95, Homer1, synaptophysin, Rab5, Rab7, and Rab11 were normal in PAC KD neurons, indicating that PAC is not essential in regulating the expression of these synaptic and endosomal proteins under basal conditions.

We then assessed whether regulation of endosomal pH by PAC is functionally relevant for AMPAR trafficking in live neurons. To address this, we used control of PAC KD neurons co-expressing superecliptic phluorin (SEP) tagged to the GluA2 AMPAR subunit (SEP-GluA2), a widely-used tool to track AMPAR dynamics at dendritic spines, and measured AMPAR trafficking dynamics during basal conditions and chemical LTP. We first employed fluorescence recovery after photobleaching (FRAP) to measure the lateral diffusion of SEP-GluA2 within dendritic spines. Spine SEP-GluA2 FRAP had an estimated time constant (τ) of 4.469 minutes for control neurons and 3.856 minutes for PAC KD neurons which were comparable to previously reported values ^30^ (Fig. 3A-B). To measure AMPAR delivery to dendritic spines during chemical LTP (cLTP), we induced cLTP with glycine treatment and measured SEP-GluA2 intensity in dendritic spines throughout a 60-minute period. As expected, spines of control neurons underwent rapid fluorescence intensity increase upon glycine stimulation, which was maintained for 50 minutes. PAC KD neurons also showed a similar increase in SEP intensity throughout the experiment compared to controls (Fig. 3C-D). Interestingly, we observed a slight increase in trafficking kinetics for PAC KD neurons in both FRAP and cLTP experiments, but these changes were not statistically significant. Overall, PAC and endosomal hyper-acidification do not appear to be involved in the lateral mobility of AMPARs in unstimulated and glycine-stimulated neurons.

**Figure 3.**
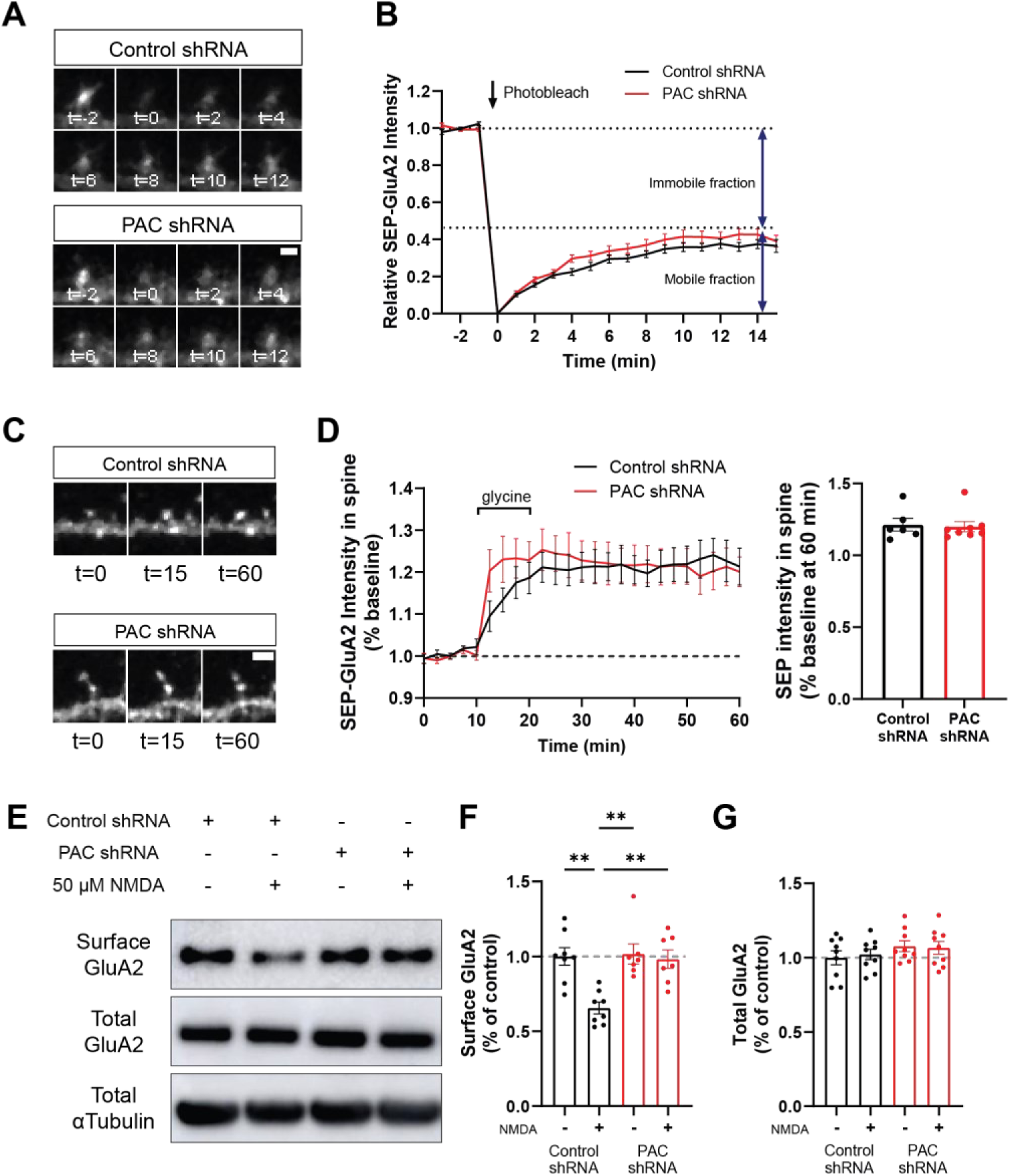
PAC is required for AMPAR internalization during LTD but does not affect LTP or basal trafficking. **(A)** Representative images of control and PAC KD neurons expressing SEP-GluA2 undergoing photobleaching. Scale bar 2 µm. **(B)** Quantification of FRAP in control (n = 25) and PAC KD (n = 29) neurons showing similar fluorescence recovery dynamics. **(C)** Representative images of control and PAC KD dendritic spines in response to glycine induced cLTP. Scale bar 2 µm. **(D)** Quantification of cLTP in control and PAC KD neurons (left) over the time course of the assay and (right) at 60 min post stimulation. SEP intensity was comparable between control and PAC KD neurons. Two-tailed student’s t-test. **(E)** Western blot depicting expected decrease in surface GluA2 following NMDA stimulation in control neurons. PAC KD neurons do not internalize surface GluA2 following NMDA. **(F and G)** Quantification of surface GluA2 and GluA2 protein levels after cLTD and surface biotinylation. Error bars represent mean ± SEM. **p < 0.01, two-way ANOVA with Tukey correction for multiple comparisons.

### Activity-dependent AMPAR internalization is impaired during chemical LTD in PAC KD neurons

We next measured AMPAR internalization during chemical LTD (cLTD) using N-Methyl-D-aspartic acid (NMDA) treatment. Due to confounding changes in intracellular pH caused by NMDA administration, SEP-GluA2 cannot faithfully measure internalization of AMPARs ^31^. Instead, we employed a surface biotinylation assay following the cLTD stimulus to compare surface (sGluA2) versus total (tGluA2) levels of the AMPAR subunit GluA2 in control and PAC KD neurons. Using this approach, we observed a ∼30% decrease in surface GluA2 expression following NMDA treatment in control shRNA-treated cortical neurons (Fig. 3E). Interestingly, PAC KD neurons failed to internalize GluA2 after NMDA stimulation (Fig. 3F). As a control, we measured total GluA2 levels (Fig. 3G), which were unchanged by NMDA treatment.

AMPARs can be endocytosed through a constitutive trafficking pathway and a regulated activity-dependent pathway that have distinct molecular mechanisms ^32–34^. LTD employs the activity-dependent pathway, which utilizes calcium-sensing proteins to increase AMPAR endocytosis. Homeostatic scaling is an alternative method of globally regulating surface AMPAR levels on a longer timescale and has been reported to use the constitutive trafficking pathway. Homeostatic downscaling of AMPARs and constitutive AMPAR trafficking was reported to use clathrin- and dynamin-independent mechanisms, whereas LTD relies on clathrin and dynamin ^34,35^. To assess homeostatic downscaling of surface AMPARs, we employed surface biotinylation following 48 hours of bicuculine treatment in neuronal cultures. We observed robust downregulation of surface GluA2 in both control and PAC KD neurons after bicuculine treatment (Fig. S4A-C), suggesting that endocytic mechanisms involved in homeostatic scaling are not dependent on the PAC channel. Thus, PAC is required for acute activity-dependent endocytosis of surface AMPAR during LTD but is not involved in mechanisms regulating constitutive trafficking or homeostatic plasticity.

### A novel live-cell imaging cLTD assay reveals impaired AMPAR endocytosis in PAC KD neurons

In addition to biochemically probing surface AMPAR content following cLTD, we developed a new imaging paradigm to detect AMPAR endocytosis in real-time in live-neurons. Previously, transient intracellular pH decreases following NMDA treatment quenched the fluorescence of internal SEP-AMPAR located in dendrite endoplasmic reticulum. To circumvent this, we expressed HaloTag-GluA2 in neurons and labeled AMPARs with a cell-impermeable, pH-sensitive Halo dye, AcidifluorORANGE, which increases fluorescence intensity as pH decreases (Fig. 4A). This paradigm avoids non-specific labeling of intracellular AMPARs and specifically labels surface and newly endocytosed AMPARs, which is measured through the appearance of fluorescent puncta as AcidifluorORANGE-labeled AMPARs accumulate in the acidified lumen of endosomes. Minimal internalization is observed in untreated cells aside from normal cycling of AMPARs through the constitutive endocytic pathway. However, NMDA treatment in rat primary hippocampal neurons induces a robust increase in internalized AMPARs throughout the dendrites and soma that persists for at least 60 minutes and mostly undergo retrograde transport (Fig. 4B, Movie S1). To validate the specificity of this new LTD assay, we applied D-APV, an NMDAR blocker that inhibits both LTP and LTD, to NMDA-treated neurons and found no increase in AMPAR endocytosis (Fig. 4C-E). We then measured AMPAR internalization in PAC KD neurons and PAC KD neurons re-expressing WT, YLAA, or K319A hPAC (Fig. 4F-H). In agreement with our surface biotinylation data, PAC KD neurons did not endocytose new AMPARs after NMDA stimulation, and endosomes remained static along dendrites (Fig. 4F-H, Movie S1). This result suggests that regulation of endosomal pH by PAC is required for AMPAR endocytosis and endosome dynamics during LTD. Overexpression of WT PAC, but not the YLAA and K319A mutants, rescued AMPAR endocytosis, suggesting that, similar to pH regulation, proper PAC endosomal localization and Cl^-^ channel function are both required for the expression of LTD.

**Figure 4.**
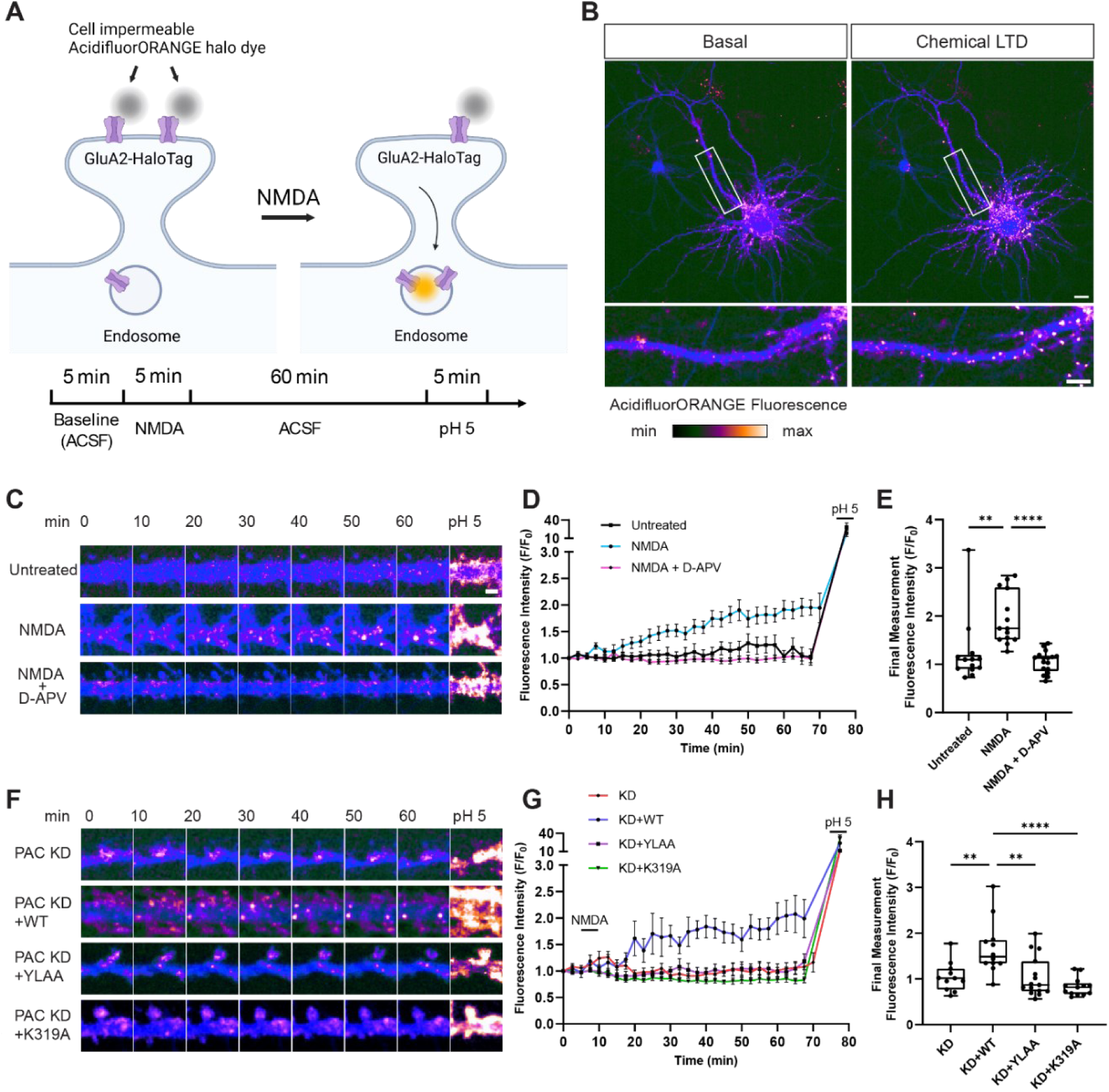
Live-cell chemical LTD assay tracks the endocytosis of GluA2, which is impaired in PAC KD neurons. **(A)** Schematic of live-cell chemical LTD imaging assay. Neurons expressing Halo-GluA2 are loaded with the pH-sensitive halo-dye AcidifluorORANGE prior to induction of cLTD using NMDA. **(B)** Representative neuron expressing Halo-GluA2 and GFP cell-fill (blue) before and after NMDA stimulation, demonstrating the internalization of HaloTagged AMPA receptors. Scale bars: upper, 10 µm; lower, 5 µm. (C) Representative images of neurons treated with NMDA (n = 14), untreated (n = 13), or treated with both NMDA and D-APV (n = 17). Scale bar 2 µm. (D) Quantification of fluorescence intensity of AcidifluorORANGE signal during the assay. (E) Quantification of final versus initial fluorescence intensity for each condition. (F) Representative images of PAC shRNA treated neurons alone (n = 11) or along with WT PAC cDNA (n = 12), or the PAC mutants YLAA (n = 15) or K319A (n = 13). (G) Quantification of fluorescence intensity of AcidifluorORANGE signal during the assay for PAC KD background neurons. (H) Quantification of final versus initial fluorescence intensity for each condition. Error bars represent mean ± SEM. *p < 0.05, **p < 0.01, ***p < 0.001, ****p < 0.0001, two-way ANOVA with Tukey correction for multiple comparisons.

### Pyramidal neuron-specific PAC KO mice have normal dendritic spine density and basal synaptic transmission

To investigate the physiological consequences of PAC deletion in neurons, we generated a conditional PAC knockout (cKO) mouse where PAC is specifically deleted in pyramidal neurons in the hippocampus and neocortex by crossing PAC^F/F^ with the NEX-Cre line ^36^. The PAC cKO mice were viable and grossly normal. Transcript levels of PAC in whole hippocampal tissue were significantly reduced, with remaining transcripts likely originated from glial cells. The dendritic spine density of hippocampal CA1 pyramidal neurons in PAC cKO mice (1.076 spines/µm) and littermate controls (0.9997 spines/ µm) were similar (Fig. S5C). To test whether PAC deletion affects excitatory synaptic transmission, we prepared acute hippocampal slices and measured miniature excitatory post-synaptic currents (mEPSCs) in CA1 neurons using whole-cell electrophysiology in the presence of TTX to block neuronal activity (Fig. S5D). The amplitude of mEPSCs, which can be attributed to post-synaptic AMPAR density or pre-synaptic vesicle loading, was similar between WT and PAC cKO mice (Fig. S5E). The frequency of mEPSCs, determined by synapse number and vesicle release probability, was also comparable between WT and PAC cKO mice (Fig. S5F). We then evoked CA1 EPSCs by stimulating the Schaffer collateral axons to determine how PAC deletion would affect synaptic transmission, but did not observe any difference in evoked EPSC amplitude between groups (Fig. S5G-H). Finally, we used voltage clamp to quantify CA1 AMPAR and NMDAR properties by measuring current amplitude at −70 mV and +40 mV, respectively. The resulting AMPA/NMDA ratio, which describes synaptic content and strength, was also unchanged in PAC cKO mice (Fig. S5I-J). Taken together, we conclude that the PAC channel is dispensable for hippocampal excitatory synaptic transmission.

### PAC cKO mice have normal hippocampal LTP, but impaired NMDAR-dependent and mGluR-dependent LTD

Based on our observation that PAC cKO mice have impaired activity-dependent AMPAR internalization, we tested if synaptic plasticity was impaired in hippocampal brain slices. We measured the Schaffer-collateral-evoked field excitatory postsynaptic potential (fEPSP) slope after inducing NMDA receptor dependent LTP and LTD with theta-burst stimulation (TBS) and low-frequency stimulation (LFS), respectively. Interestingly, PAC cKO CA1 hippocampus exhibited markedly impaired LTD, while LTP remained normal (Fig. 5A, B). To examine the role of PAC in synaptic plasticity at single-cell levels, we performed whole-cell recordings in hippocampal CA1 pyramidal neurons. Consistent with the fEPSP measurements, PAC cKO neurons exhibited normal LTP but impaired LTD (Fig. 5C, D). Additionally, treatment of hippocampal slices with dihydroxyphenylglycine (DHPG) to activate metabotropic glutamate receptor (mGluR)-dependent LTD also induced robust LTD in WT but not in PAC cKO neurons (Fig. 5E), suggesting that PAC deletion impairs these distinct LTD mechanisms through AMPAR internalization, which occurs downstream of both forms of LTD ^37^. To assess whether the LTD defect is a cell-autonomous phenotype, we sparsely expressed hSyn-Cre-mCherry in CA1 neurons from PAC^F/F^ mice using *in utero* electroporation. EPSCs were measured from neurons originating from the same mouse to compare LFS-induced LTD by whole-cell recordings. Cre-mCherry-negative control neurons had robust LTD whereas Cre-mCherry-positive neurons had impaired LTD (Fig. 5F). Given the sparse transfection of the plasmid, we conclude that defects in LTD are cell autonomous and are likely caused by impaired AMPAR endocytosis rather than changes in presynaptic release. Taken together, our results suggest that the regulation of endosomal pH through PAC is required for NMDAR- and mGluR-dependent LTD in the hippocampus but is not essential for LTP.

**Figure 5.**
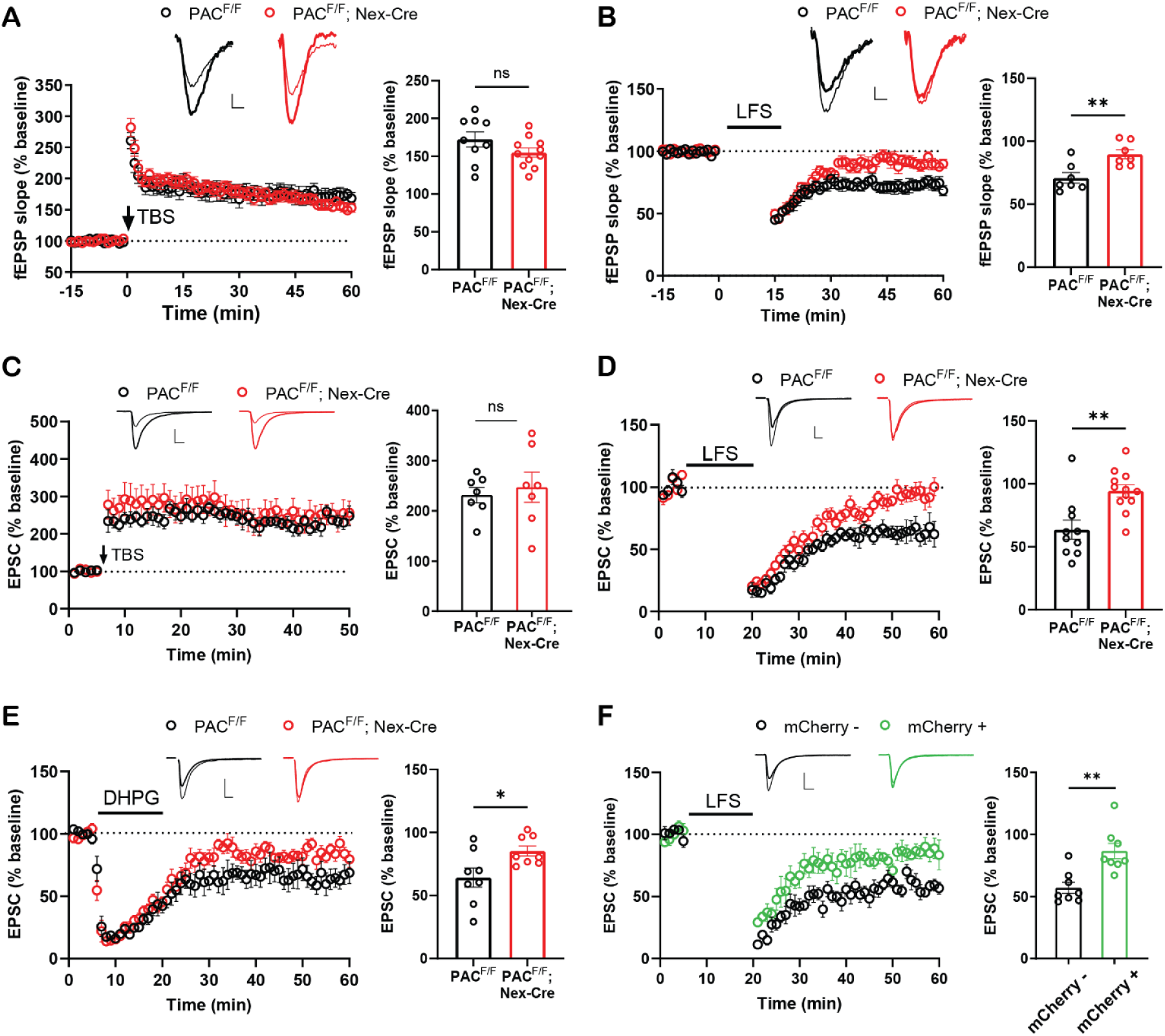
PAC cKO mice have impaired NMDAR- and mGluR-dependent LTD, but normal LTP. Field recordings from acute hippocampal slices: **(A)** TBS-induced LTP at the Schaffer collateral-CA1 synapses from PAC cKO mice is comparable to wild-type. **(B)** LFS-induced LTD is impaired in PAC cKO mice. Whole-cell recordings from CA1 pyramidal neurons: **(C)** TBS-induced LTP is normal in PAC cKO neurons. **(D)** LFS-induced NMDAR-dependent LTD is impaired in PAC cKO neurons. **(E)** DHPG-induced mGluR-dependent LTD in PAC cKO neurons. **(F)** LFS-induced LTD in sparsely electroporated CA1 neurons with expressing hSyn-Cre-mCherry is impaired compared to Cre-mCherry-negative neurons from the same mice. Quantification of fEPSP slopes or EPSCs for the final 10 minutes of the protocol are quantified on the right for each panel. (A-F) Sample traces represent fEPSPs (A-B) or EPSCs (C-F) at baseline (thin) and 1 hr after (thick) stimulation. (A-B) Scale bars represent 0.1 mV (vertical), 5 ms (horizontal). (C-F) scale bars represent 50 pA (vertical), 10 ms (horizontal). Recordings were conducted on male and female 1-2-month-old mice. Data are reported as mean ± SEM. **p < 0.01, *p<0.05, ns: not significant. Student’s t-test.

### PAC cKO mice have impaired reversal learning

Impairments in synaptic plasticity often manifest in cognitive and behavioral deficits. We conducted a battery of behavioral tests in male PAC cKO mice alongside WT littermate controls to assess cognitive and behavioral function. PAC cKO mice displayed normal locomotion during 30 minutes of testing in the open field test (Fig. S7A). Anxiety, measured in the elevated plus maze, was comparable between WT and cKO mice (Fig. S7B). We next performed behavioral assays that require functional LTP, including the Y-maze, contextual fear conditioning, and the Morris Water Maze (MWM). Since hippocampal LTP appears normal in PAC cKO mice, we added an additional reversal training session and probe to the MWM assay to assess LTD. As expected, assays dependent on hippocampal LTP, including the spontaneous alternation in the Y-maze (Fig. S7C), contextual fear memory following fear conditioning (Fig. S7D-F), and MWM spatial acquisition (Fig. 6A, B) were all normal in PAC cKO mice. The MWM reversal task is dependent on hippocampal LTD for behavioral flexibility, or the ability to appropriately adapt to a changing environment ^14,38,39^. Interestingly, cKO mice performed worse in the reversal task during the first two days of reversal training (Fig. 6A). PAC cKO mice also performed worse than WT mice during probe trial 2 (Fig. 6C), spending reduced time in the correct target quadrant. Swimming speeds were comparable between WT and cKO mice (Fig. 6D). Altogether, our behavioral results are consistent with our *in vitro* and *ex vivo* data, demonstrating that PAC deletion in pyramidal neurons impairs LTD, while basal synaptic transmission and LTP are maintained. This is reflected in the behavior of PAC cKO mice, where otherwise normally performing mice have impaired behavioral flexibility.

**Figure 6.**
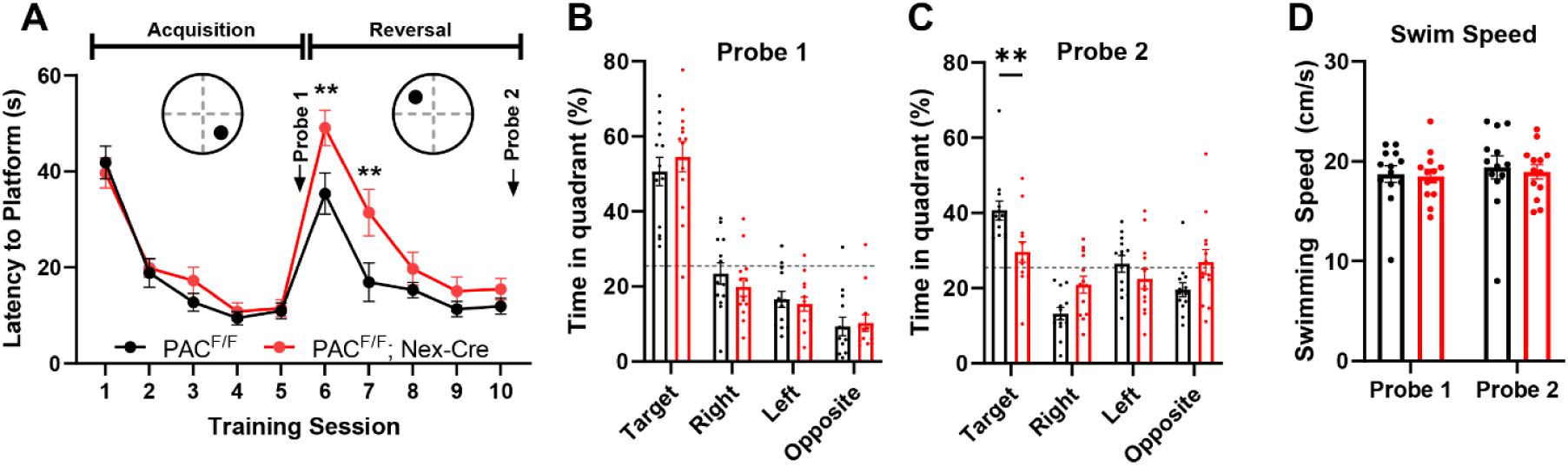
PAC cKO mice have impaired reversal learning. **(A)** Latency to platform during the spatial acquisition phase and the reversal phase during Morris Water Maze training. Escape latencies were similar for WT and PAC cKO mice during the acquisition phase, but cKO mice have delayed latencies during the first two days of reversal training (training sessions 6 and 7). **(B)** Quantification of time spent in each quadrant during probe trial 1, following acquisition training. **(C)** Quantification of time spent in each quadrant during probe trial 2, following reversal training. PAC cKO mice spend significantly less time in the correct target quadrant compared to WT controls. **(D)** Quantification of swim speeds for WT and cKO mice. Male mice aged 2-5 months were used for the behavioral assay (WT, n = 13; cKO n = 13). Data represent mean ± SEM **p < 0.01. Two-way ANOVA with Bonferroni’s correction for multiple comparisons.

## Discussion

In this study, we utilized HaloTag-GluA2 as a new approach to track AMPAR endocytosis in live neurons and revealed a prominent role of the PAC channel and endosomal hyper-acidification in regulating activity-dependent AMPAR internalization and LTD in hippocampal neurons. While both LTP and LTD are important for synaptic plasticity, the molecular tools to study LTD are less developed than those for LTP, and many inhibitors of LTD also affect LTP ^37^. LTD is essential for cognitive functions such as memory, behavioral flexibility, and novelty detection ^37^. Furthermore, dysregulation of synaptic plasticity is implicated in neurodegenerative diseases such as Alzheimer’s disease (AD), where aberrant cellular signaling may cause pathological synaptic depression and loss ^40–42^. Defects in endosomal acid-base regulation have also been implicated in AD ^22^. Thus, targeting PAC to restore endosomal function may offer a potential therapeutic strategy for mitigating deficits in synaptic plasticity.

It is well established that proper pH or Cl^-^ regulation within the endolysosomal system is crucial for neural development and synaptic function, as evidenced by studies on endosomal sodium proton exchangers NHE6 and NHE9 ^22^ or chloride transporters CLC-3 ^43,44^ and CLC-4 ^45–47^. Loss-of-function of NHE6/9 results in hyper-acidified endosomes and outcomes range from impaired neurite development ^20^, dendritic spine density ^48^, endolysosomal maturation ^21^, and synaptic transmission ^19^. Similarly, endosome alkalinization also results in impaired neural phenotypes, most notably abolishing LTD and AMPAR endocytosis in the cerebellum ^10^ and disrupting dendritic branching and cargo uptake ^49^. Neuronal PAC functions as an early and recycling endosome channel that regulates luminal pH, which is consistent with other reports describing the subcellular localization of PAC and its role in preventing luminal hyper-acidification ^24,25^. While PAC is also present on the plasma membrane, our data indicate that the main function of PAC is endosomal. Based on our localization and electrophysiology data, it seems unlikely that PAC regulates pre-synaptic plasticity. Interestingly, the consequences of PAC deletion are more specific and less severe compared to the deletions of NHE6, NHE9, or CLC-3, which cause neurodevelopmental defects. Instead, PAC is directly involved in LTD by facilitating the rapid removal of AMPARs from the synapse through endocytosis.

Acute regulation of endosomal pH, whether through optogenetic alkalinization ^10^ or through PAC deletion, directly inhibits activity-dependent AMPAR endocytosis. To our knowledge, these are the only examples directly linking altered endosomal pH with an AMPAR trafficking defect during LTD. Interestingly, LTP is normal in mice with either over-alkalinized or over-acidified endosomes, supporting a model where LTP is less reliant on endocytic machinery for the early stages of AMPAR delivery to the PSD. Indeed, the lateral diffusion of AMPARs is a major contributor to receptor enrichment at dendritic spines during LTP ^3,4,50^. Our data support this model, but it remains possible that impaired endosomal acidification may be involved in later stages of LTP, where AMPARs need to be exocytosed from intracellular stores to replenish membrane AMPAR levels ^51^. Currently, we can only speculate on the mechanisms by which alkalinization or hyper-acidification affects AMPAR endocytosis. Proteins involved in AMPAR trafficking such as PICK1, Rab5, Rab7, or Rab11 and their effectors are all candidates for proteins that can be disrupted by altered pH.

To directly observe AMPAR endocytosis in live neurons, we developed a cLTD assay utilizing HaloTag-GluA2 and the AcidifluorORANGE Halo-dye. This assay addresses a known limitation of SEP tagged AMPARs where NMDA causes nonspecific fluorescence quenching of intracellular SEP-AMPARs located in the endoplasmic reticulum ^31^. Using HaloTag-GluA2 provides several advantages. First, a variety of Halo dyes with different properties can be used to label AMPARs in a time-controlled manner. We are able to measure newly endocytosed AMPARs by using a membrane impermeable dye that becomes fluorescent in the low pH environment of the endosome. Second, this method allows for the longitudinal tracking of endocytosed AMPARs in neurons over the whole period of LTD induction, allowing the direct visualization of endocytic events, endosomal maturation and fusion, and anterograde or retrograde trafficking, which have not been previously observed. In untreated neurons, our assay reveals both endocytic and exocytic events, though there is rarely a net increase in endocytosed AMPARs over time. During cLTD, we observed robust AMPAR endocytosis, maturation, and retrograde transport, which were absent in PAC KD neurons. The main limitation of this assay is that internalized receptors can only be visualized after luminal acidification such that the earlier stages of endocytosis may not be detectable. Despite this, measuring receptor trafficking dynamics with our cLTD assay may prove valuable for future studies investigating the mechanisms underlying the complex cellular trafficking events of AMPARs and can be adapted for *in vivo* experiments.

PAC deletion specifically disrupts LTD but does not affect LTP or basal synaptic transmission. This phenotype mirrors that of mice lacking calcium-sensitive proteins PICK1 ^11,12^, which is involved in the endocytosis and retention of GluA2, and synaptotagmin-3 ^14^, which is crucial for the activity-dependent endocytosis of GluA2. Our results are consistent with a model in which AMPARs can be endocytosed through two distinct molecular pathways: a constitutive trafficking pathway that maintains surface AMPAR content and an activity-dependent regulated pathway that reduces surface AMPAR levels ^32–35^. The regulated pathway involves the specific removal of GluA2-containing AMPARs from the membrane following PKC phosphorylation at the S880 residue ^2,52^. GluA2 first dissociates from the PSD and diffuses to endocytic zones. Endocytosis occurs either through direct binding to the AP-2 adaptor complex ^53,54^ or via AMPAR-interacting proteins that deliver AMPARs to the endocytic machinery. In contrast, slower endocytic events such as constitutive trafficking and homeostatic downscaling utilize a clathrin-independent pathway ^35^. The presence of these two distinct pathways may explain why only the activity-dependent pathway is impaired in our model. Based on our data and the existing literature, we conclude that PAC and endosomal acidification are required only for rapid, activity-dependent AMPAR endocytosis, whereas the constitutive pathway appears to be less sensitive to perturbations in endosomal pH.

We have shown that PAC is important for endosomal pH balance and LTD in neurons. However, the molecular players involved in linking endosomal pH to LTD remain elusive due to redundancy in endocytic machinery ^55,56^. Nevertheless, the discovery that endosomal hyper-acidification ablates hippocampal LTD serves as a counterpart to the previous finding that endosomal alkalinization impairs LTD in the cerebellum ^10^ and has broader implications in neurobiology. Currently, it is unknown whether the LTD impairments caused by hyper-acidification and alkalinization of the endosome result from the same or distinct mechanisms. It would be important to test whether trafficking of other neurotransmitter receptors is similarly affected by altered endosomal acidification. Additionally, exploring whether PAC is involved in LTD and AMPAR trafficking in Purkinje cells of the cerebellum, and its impact on motor learning, could provide further insights. Dysregulation of endosome function is also implicated in autism and AD, but it remains unclear if PAC contributes to the pathogenesis of these disorders. As proper LTD and endosomal function is important for synaptic function throughout the CNS, it is likely that future studies will uncover novel roles of PAC in different physiological and pathological contexts.

## Materials and Methods

### Animals

All procedures related to animal care and treatment conformed to Johns Hopkins University Animal Care and Use Committee guidelines. Animals were group housed in a standard 12-hour light/dark cycle with *ad* libitum access to food and water. Male and female animals were used for all experiments unless otherwise stated. The *NEX-Cre* line was obtained as a gift from Klaus-Armin Nave ^36^. *PAC^F/F^* mice were generated previously^57^ and crossed with the *NEX-Cre* line to generate *PAC^F/F^; NEX-Cre* PAC cKO mice. Sprague Dawley rats (Harlan Laboratories) were used for hippocampal or cortical cultures at embryonic day 18 (E18) as described below. All animals were group housed in a standard 12 hr light/ 12 hr dark cycle with *ad libitum* access to food and water.

### DNA Constructs

pFSW-hSyn-mVenus, SEP-GluA2, and HaloTag-GluA2 plasmids were gifts provided by Richard Huganir. hPAC overexpression constructs were generated by cloning hPAC into lenti-EGFP-Cre (a gift from Shuying Sun) downstream of the EF-1α core promoter. hPAC fusion constructs were cloned from the overexpression construct by inserting a linker followed by either EGFP or mCherry at the 3’ end of PAC cDNA. Plasmid vectors (pLL3.7) containing control and *rat* PAC shRNA were adapted from Osei-Owusu et al. ^28^. pLL3.7 shRNA vectors expressing mCherry were cloned from the original vectors. Rescue vectors were cloned by replacing EGFP with hPAC-P2A-EGFP, which is resistant to *rat* PAC shRNA. Vectors expressing hPAC Y10A/L13A or K319A mutant cDNA were cloned from the WT hPAC rescue vector using site-directed mutagenesis. Transferrin receptor conjugated ratiometric pHluorin (TfR-RpH) was generated by cloning ratiometric pHluorin (Addgene 163070) into pcDNA3-hTfR-pHuji (Addgene 61505). The pFSW-hSyn-Cre-P2A-mCherry vector used for in utero electroporation was generated by cloning Cre-P2A-mCherry into the pFSW-hSyn-mVenus vector.

### Primary Neuron Cultures and Transfection

Primary rat neuron cultures were performed as previously described with some modifications ^15^. Hippocampal or cortical neurons dissected from E18 rat pups were plated onto poly-D-lysine coated coverslips or plates in NM5 (Neurobasal medium with 5% horse serum, 2% B27, 2 mM Glutamax, 50 U/ml penicillin and 50 mg/streptomycin). Hippocampal neurons were switched to NM0 (Neurobasal medium 2% B27, 1 mM Glutamax, 50 U/ml penicillin and 50 mg/streptomycin) one day post-seeding and fed once a week with NM0. Cortical neurons were fed at day-in-vitro (DIV) 5 with NM0 plus FDU (5 mM 5-Fluro-2′-deoxyuridine and 5 mM Uridine) to stop glia proliferation, then fed twice a week with NM0 without FDU. For staining and live imaging, hippocampal neurons were plated at a density of approximately 40,000 cells/cm^2^. Neurons were transfected at DIV11-16 using Lipofectamine 2000 (Invitrogen) following the manufacturers protocol, and the cells were used 3-5 days later. For biochemistry, cortical neurons were plated at a high density of 100,000 cells/cm^2^ into 6-well plates and were used at 2-3 weeks old. Primary mouse hippocampal neurons were obtained from P0 mouse pups using the same procedure and culturing conditions as rat neurons.

### Human iPSC-derived Neurons

Human-iPSCs were cultured based on a previously established protocol with slight modifications ^58^. Human-iPSCs were grown on Matrigel coated plates in Essential 8 basal medium. For stage 1 of differentiation, iPSCs were seeded in N2 medium (DMEM-F12 containing 1X N2, NEAA, and GlutaMax, 2 ug/ml Doxycycline, and 10 mM Rock inhibitor) and allowed to differentiate for 3 days with daily media changes. For stage 2 of differentiation, iPSCs were dissociated with accutase and plated onto PLO coated coverslips in complete Brianphys media (Brainphys with B-27, 10 ug/ml BDNF, 10 ug/ml NT3, 1 mg/ml Laminin, and 2 ug/ml Doxycycline). Half medium changes were performed every other day and cells were used after 14 days of differentiation.

### Lentivirus Production and Transduction

Active lentiviral particles were generated using the 3^rd^ generation lentivirus system, consisting of pRRE, pREV, and pVSVG. Packaging vectors were co-transfeted with specific lentiviral vectors into 80-90% confluent HEK293T cells using PeneFect transfection reagent according to the manufacturers protocol. One day post-transfection, the media was replaced with DMEM supplemented with 10% FBS, 1% BSA, and 1% penicillin/streptomycin. The supernatant was collected 48-72 hours after the media change and filtered through a PDVF membrane with 0.45 um pore size. Lentivirus was concentrated using LentiX concentrator according to the manufacturers protocol and resuspended in Neurobasal medium. Lentivirus was either used immediately or kept at – 80°C for long-term storage. All reagents that came into contact with lentivirus were disinfected with bleach for 24 hours before disposal. The viral titer was calculated before lentiviral transduction into cultured neurons. Transduction was performed by incubating the lentivirus with neuronal cultures overnight. 100% of the media was removed the next day and replaced with 50% conditioned media and 50% fresh NM0. Cultures were used for experiments 96 hours post-transduction.

### Immunocytochemistry

Neuronal cultures were fixed in 4% paraformaldehyde (PFA) and 4% sucrose (w/v) in PBS, then permeabilized with 0.1% TritionX-100 in PBS for 10 minutes, followed by a 30-minute block in 3% BSA in PBS. Primary antibodies were diluted in blocking buffer and incubated overnight at 4C. Primary antibodies: chicken MAP2 (Biosensis C-1382-50, 1:2000), mouse hPAC (1:1000), rabbit EEA1 (Cell Signaling Technology C45B10, 1:1000), rabbit Syntaxin13 (Synaptic Systems 110-132, 1:500), rabbit GluA1 (JH4294, 1:1000), mouse GluA2 (15F1, 1:5000), guinea pig Homer1 (Syhnaptic Systems 160-004, 1:500), rabbit Synapsin1 (Synaptic Systems 106-103, 1:500), guinea pig Vglut1 (Synaptic Systems 135-304, 1:500). Cells were washed with PBS and incubated with secondary antibodies conjugated to fluorescent dyes (Thermo, 1:1000) for 1 hour at room temperature before final washes and mounting. Samples were imaged using a Zeiss LSM900 confocal microscope. The ImageJ plugin JaCoP was used to measure the Pearson’s R correlation coefficient from co-immunolabeled images where ROIs were selected based on the fluorescence of PAC puncta.

### Live-cell imaging for endosomal pH

Primary hippocampal neurons were co-transfected with TfR-RpH and shRNA-mCherry constructs in a 1:1 ratio at DIV11 and imaged 3-5 days later. Neurons were imaged in a custom-made live imaging chamber containing ACSF at room temperature (120 mM NaCl, 5 mM KCl, 2 mM CaCl_2_, 1 mM MgCl_2_, 10 mM glucose, 10 mM HEPES, pH 7.3) to acquire basal endosomal fluorescence at 405 nm and 488 nm excitation. The chamber solution was switched to pH calibration buffers equilibrated to pH 7.5, 6.5, and 5.5. Calibration solutions were adapted from Pohlkamp et al. ^59^. pH 7.5 buffer: for pH < 7.

Fluorescence images were acquired for both 405 nm and 488 nm stimulation for all pH solutions and were used to generate a standard curve for each experiment. Values of the 405/488 fluorescence intensity ratio from the basal ACSF condition were plotted on the standard curve to vesicular calculate pH. Imaging was performed on a Zeiss LSM900 confocal microscope.

### Live-cell imaging of SEP-GluA2 for AMPAR trafficking and chemical LTP

Primary hippocampal neurons were co-transfected with SEP-GluA2 and shRNA-mCherry constructs in a 1:1 ratio at DIV 15 and imaged at DIV 18-21 for measurements of basal AMPAR trafficking and cLTP. Live-neurons were imaged in a chamber containing ACSF at room temperature. For basal AMPAR trafficking, 2-5 dendritic spines per neuron were selected for SEP fluorescence measurement. Z-stacks were collected once per minute over a 15-minute experiment measuring fluorescence intensity. Glycine-induced chemical LTP was performed as previously described with minor modifications ^15^. Briefly, cells were pre-incubated for 1-2 hours in basal ACSF (120 mM NaCl, 5 mM KCl, 2 mM CaCl_2_, 1 mM MgCl_2_, 10 mM glucose, 10 mM HEPES, pH 7.3) supplemented with 500 nM TTX, 20 µM bicuculine, and 1 µM strychnine. Baseline was collected in basal ACSF, followed by a 10 minutes cLTP stimulation with 200 uM glycine in the same ACSF solution but without MgCl_2_. Basal ACSF was used after stimulation for the remainder of imaging (50 minutes).

### Live-cell imaging of HaloTag-GluA2 for chemical LTD

Primary hippocampal neurons were co-transfected with HaloTag-GluA2 and shRNA-EGFP constructs in a 1:1 ratio at DIV 15 and imaged at DIV 18-21. Prior to imaging, cultures were pre-incubated in basal ACSF supplemented with 1 µM TTX for 1 hour. A stock of 1mM Halo-Tag AcidifluorORANGE (Millipore Sigma, SCT212) was diluted 1:4000 in basal ACSF to for dye loading at 37 C for 15 minutes. Prior to imaging, AcidifluorORANGE was washed out with three washes of basal ACSF and transferred to a live-imaging chamber maintained at 37 C and 5% CO2. For cLTD assay, neurons were stimulated for 5 minutes with 20 µM NMDA in ACSF with 1 µM TTX and without MgCl_2_, then replaced with basal ACSF for the remainder of imaging (60 minutes). At the end of each experiment, the solution was changed to pH 5 calibration buffer (125 mM KCl, 25 mM NaCl, 10 μM monensin, 25mM MES) to confirm AcidifluorORANGE loading. Experiments performed under D-APV used the same experimental conditions but in the presence of 50 μM D-APV. For no-stimulation experiments, neurons were imaged for 60 minutes in basal ACSF, followed by pH 5 calibration buffer. Images were obtained on either Nikon Eclipse Ti2 or Zeiss 3i spinning disc microscopes with a 50 µm pinhole size. Images were processed in ImageJ-FIJI. To quantify fluorescence intensity of endocytosed AcidifluorORANGE, images were thresholded and the particle analysis function was used to obtain particle number, area, and integrated density. Integrated fluorescence intensities were plotted against baseline values to obtain fluorescence change curves (F/F_0_).

### Chemical LTD and Surface Biotinylation

To induce chemical LTD, cultured hippocampal or cortical neurons were incubated with 50 uM NMDA for 2 minutes at 37C in culturing medium. Afterwards, neurons were incubated in NMDA-free culturing medium for 15 minutes at 37C. After cLTD, neurons were placed on ice for 5-10 minutes. Surface biotinylation was performed as previously described with some modifications ^60,61^. Briefly, neurons were rinsed once with ice-cold PBSCM [PBS-calcium-magnesium: 1 × PBS, 1mM MgCl2, 0.1mM CaCl2 (pH 8.0)] were incubated with Sulfo-NHS-SS-biotin (1 mg/mL, Thermo Scientific) for 20 min at 4 °C. Neurons were then washed with PBSCM and incubated in 20 mM glycine twice for 5 min to quench unreacted biotinylation reagent. Neurons were lysed in lysis buffer [PBS containing 50 mM NaF, 5mM sodium pyrophosphate, 1% Nonidet P-40, 1% sodium deoxycholate, 0.02% SDS, and protease inhibitor mixture (Roche)]. Equal amounts of proteins were incubated overnight at 4 °C with NeutrAvidin agarose beads (Thermo Scientific) and then washed with lysis buffer three times. Biotinylated proteins were eluted using 2× SDS loading buffer. Surface or total proteins were then subjected to SDS/PAGE and analyzed by Western blot.

### Antibodies

The following primary antibodies were used for Western blotting: mouse alpha tubulin (Invitrogen DM1A, 1:2000), mouse GluA2 (Millipore MAB397, 1:2000), mouse Na/K-ATPase (Santa Cruz sc-48345, 1:2000), mouse PSD-95 (NueroMab K28/43, 1:1000), rabbit PAC (JH7916, 1:200), guinea pig Homer1 (Synaptic Systems 160-004, 1:500), rabbit synaptophysin1 (Synaptic Systems 101-011, 1:500), mouse Rab5 (Cell Signaling Technology C8B1, 1:1000), mouse Rab7 (Cell Signaling Technology D95F2, 1:1000), mouse Rab11 (Cell Signaling Technology D4F5, 1:2000).

### In-utero electroporation

E15.5 pregnant PAC^F/F^ mice were anesthetized with 1% sodium pentobarbital (0.1 mL/10g body weight) via intraperitoneal injection; then the mice were subjected to surgical procedures to expose the uterus. Each embryo was injected with 1-2 μL of hSyn-Cre expressing plasmid DNA mixed with Fast-Green into the lateral ventricle via a glass micropipette. Then, each embryo was electroporated with five 40-V pulses of 50 ms, delivered at 1 Hz, using platinum tweezertrodes in a square-wave pulse generator (BTX, Harvard Apparatus). After that, the embryos were placed back into the abdominal cavity, and the muscle and skin were sutured. Mice were placed on a 37℃ heat pad and monitored until fully awake.

### Electrophysiological recordings

Electrophysiological recordings were performed as described previously ^57^. Transverse hippocampal sections (300 um) were cut from mice (P20-P30) using a vibratome (VT-1200S, Leica) in ice-cold choline-based cutting solution containing (in mM): 110 choline chloride, 7 MgCl2, 2.5 KCl, 0.5 CaCl2, 1.3 NaH2PO4, 25 NaHCO3, 20 glucose, saturated with 95% O2 and 5% CO2. Slices were transferred to artificial cerebrospinal fluid (aCSF) containing (in mM): 125 NaCl, 2.5 KCl, 2.5 CaCl2, 1.3 MgCl2, 1.3 NaH2PO4, 26 NaHCO3, 10 glucose, saturated with 95% O2 and 5% CO2. The slices were allowed to recover for 1 h at 32°C and then at RT for at least 1 h before recoding. All recordings were made at RT in a submerged recording chamber with constant perfusion of aCSF containing 0.1 mM picrotoxin and 0.01 mM bicuculline, saturated with 95% O2/5% CO2. Whole-cell recordings from CA1 pyramidal neurons were visualized under an upright microscope (BX51WI, Olympus) with infrared optics. Synaptic responses were recorded by 3- to 5-MΩ borosilicate glass pipettes filled with intracellular solution containing (in mM): 125 K-gluconate, 15 KCl, 10 HEPES, 1 MgCl2, 4 Mg-ATP, 0.3 Na3-GTP, 10 phosphocreatine, and 0.2 EGTA (pH 7.2, osmolality 290-300 mOsm/kg). Data was acquired with pClamp 10.7 software (Molecular Device), filtered at 1 kHz and digitized at 10 kHz. In all experiments, the series resistance (Rs) was monitored throughout the recording and controlled below 20 MΩ with no compensation. Data was discarded when the series resistance varied by ≥ 20%.

Miniature EPSCs (mEPSCs) were measured in the presence of 1 mM TTX and a holding potential of −70 mV. Evoked EPSCs were measured following stimulation of Schaffer collaterals with a bipolar stimulation electrode placed in stratum radiatum at the CA1 region. LTP was induced by theta burst stimulation (TBS) consisting of a single train of 5 bursts at 5 Hz, and each burst contained 4 pulses at 100 Hz. NMDAR-dependent LTD was induced by low-frequency stimulation (1 Hz, 900 s) with a holding potentiation of −40 mV, after 5- to 10-min baseline recordings. DHPG-LTD was induced by the bath application of 100 um DHPG for 10 min.

To measure PAC current, whole-cell recordings in hippocampal neurons transfected with PAC shRNAs was conducted as described previously ^23^. The extracellular solution contained (in mM) 145 NaCl, 2 KCl, 2 MgCl2, 1.5 CaCl2, 10 HEPES, 10 glucose (300 mOsm/kg; pH 7.3 with NaOH). To activate PAC currents, a pH 4.6 solution was made with the same ionic composition without HEPES but with 5 mM Na_3_-citrate as the buffer, and pH was adjusted using citric acid. 1 µM tetrodotoxin (TTX,) and 100 µM amiloride were added to the bath to block voltage-gated sodium channels and ASIC channels, respectively. The internal solution contained (in mM) 120 CsCl, 20 TEA-Cl, 4 MgATP, 0.3 GTP, 10 HEPES, 4 QX-314, 0.5 EGTA (290-300 mOsm/kg; pH 7.2 with CsOH). Cells were held at 0 mV and voltage ramps (minimal interval, 200 ms duration) were applied from – 100 to +100 mV in 20 mV increments.

### Behavioral Analysis

Open field test was conducted as previously described ^57^. Locomotor activity was examined in adult (3–5-month-old) male mice by experimenters blinded to genotypes. The animals were placed in a chamber (18” × 18”) (Photobeam activity system-San Diego Instruments) and monitored for movement by using horizontal and vertical photobeams. Beam breaks were converted to directionally specific movements and summated at 5-min intervals over 30 min. Ambulatory activity was measured as total horizontal photobeam breaks, rearing was evaluated as total vertical beam breaks.

Anxiety was assessed using an elevated plus maze (66 cm long and 5 cm wide; San Diego Instruments Inc) as described previously ^62^. The maze consisted of two closed arms and two open arms suspended 54 cm above the ground. Immediately before testing, animals were placed, individually, into a clean cage for 5 min. Animals were placed onto the center of the elevated plus maze facing an open arm and allowed to explore for 5 min. Animal position was tracked using ANYmaze software (SD instruments).

Spatial short-term memory was tested using the Y-maze spontaneous alternation test (each arm 38 cm long, SD instruments). Mice were allowed to explore the maze for 5 minutes and were tracked with ANYmaze software. Spontaneous alternation was detected and calculated using ANYmaze software.

Morris water maze test was performed as previously described ^57^ with additional reversal testing. The arena consisted of a circular pool (diameter of 120 cm) filled with water that was at 24°C and made opaque with non-toxic white tempera paint. A circular, plexiglass platform (length of 10 cm) was submerged 1 cm below the surface of the water and four local cues were provided to allow spatial map generation. Mice were trained for a total of 20 trials over 5 days, with 4 trials per session and 1 session per day. Prior to the first training trial, mice were given a single habituation trial without the platform to assess any spatial bias. Trials were 60 s and mice that did not find the platform within that time were guided to the platform by the experimenter. Once on top of the platform, mice were left for an additional 10 s before being removed. Start locations (north, south, east, and west) were pseudo-randomized so that each start location was used once per session and the sequence of start locations in any session was never used twice. A probe trial was performed 24 hours after the last training. Reversal training was conducted during the next 5 days with the same parameters as the acquisition training, except that the platform was placed in the opposite quadrant. A second probe trial was performed 24 hours after the last reversal training. During probe trials, the platform was removed, and the mouse was allowed to swim freely for 60 seconds. Tracking and analysis of animal movement was done using the ANY-maze tracking system (SD instruments). Data were analyzed by comparing quadrant preferences and escape latencies averaged across animals within groups (control or cKO).

## Acknowledgments

The authors thank Seth Margolis for insightful discussion; Andrew Holland for providing microscope access; and Marie Hardwick for NEX-cre mice from Klaus-Armin Nave’s laboratory. This work was supported by grants from the National Institute of Health (R01NS118014, R35GM124824, and RF1NS134549 to Z.Q and R37NS036715 to R.L.H). Z.Q. was also supported by the American Heart Association (AHA) Established Investigator Award, McKnight Scholar Award, Klingenstein-Simon Scholar Award, Sloan Research Fellowship in Neuroscience, and Randall Reed Scholar Award. Research reported in this publication was supported by the Office of the Director of the National Institutes of Health under award number S10OD034393 to Scot Kuo for use of the Johns Hopkins Microscope Facility. J.Y. was supported by the AHA postdoctoral fellowship 20POST35200185 and Career Development Award 857671. Biorender was used to generate several figures.

