## Supporting Figures for "Endosomal hyper-acidification via proton-activated chloride channel deletion in neurons impairs AMPA receptor endocytosis and LTD"

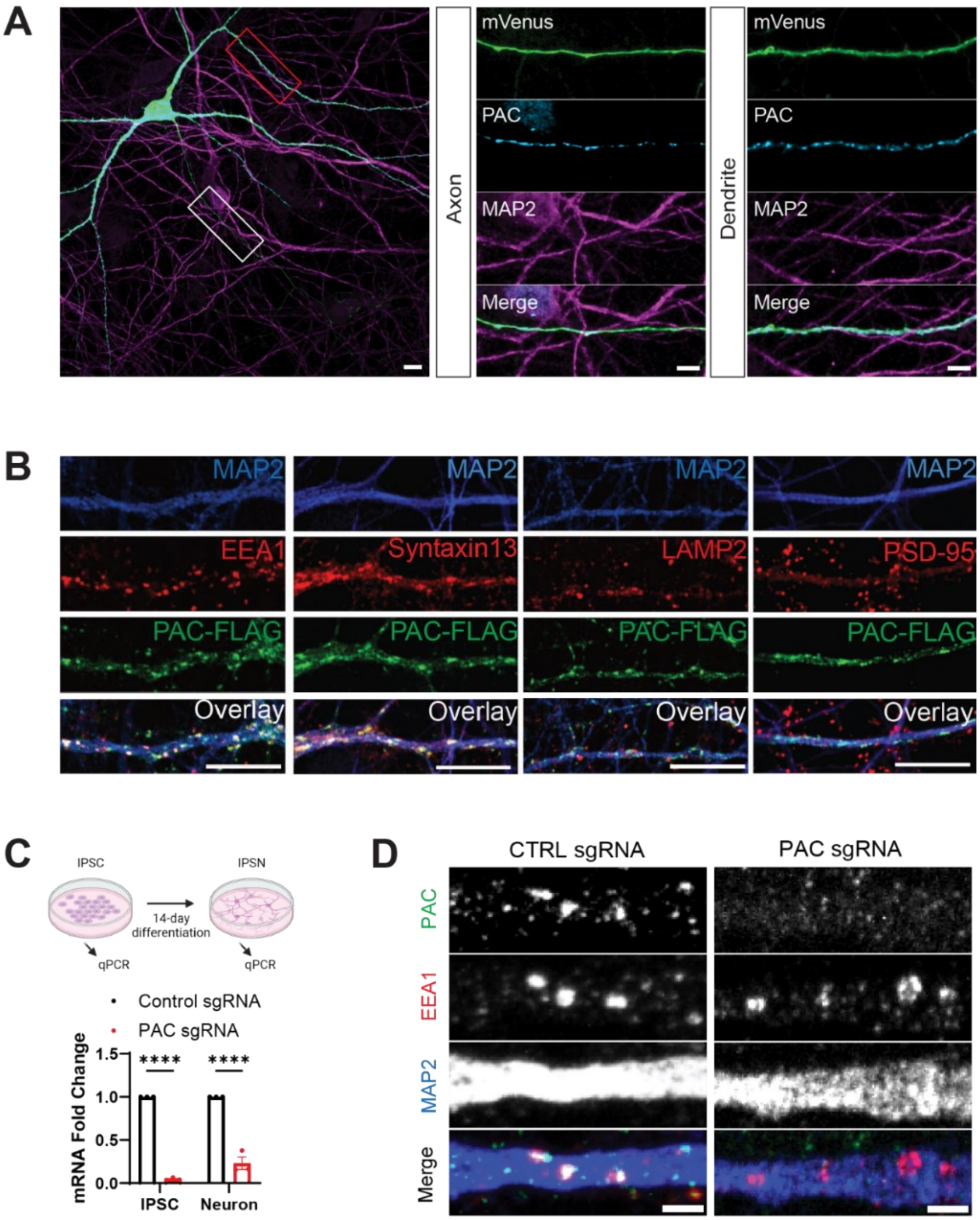


**Fig. S1. PAC localizes to dendritic endosomes in mouse primary hippocampal neurons and in human IPSC-differentiated neurons.**

**(A)** (Left) Representative image of an immunolabeled rat primary hippocampal neuron expressing hPAC and cytosolic mVenus. Scale bar 10 μm. (Middle, enlarged from white outline) MAP2-negative axon from and (Right, enlarged from red outline) MAP2-positive dendrite expressing PAC with punctate morphology. Scale bars 2 µm. **(B)** Representative images of mouse primary hippocampal neurons expressing PAC. Immunofluorescence labeling reveals co-localization of PAC and early endosomes (EEA1) and recycling endosomes (syntaxin13). PAC does not strongly co-localize with lysosomes (LAMP2) or the post-synaptic density (PSD95). Scale bar 10 µm. **(C)** Scheme for differentiating human IPSCs into neurons and qPCR validation of CRISPRi-mediated knockdown of PAC. **(D)** Representative images of human IPSC-derived neurons (IPSNs) treated with either control or PAC sgRNAs. Immunofluorescence labeling reveals endogenous PAC colocalizes with dendritic early endosomes (EEA1). PAC signal is absent from PAC KD neurons. Scale bar 2 µm.

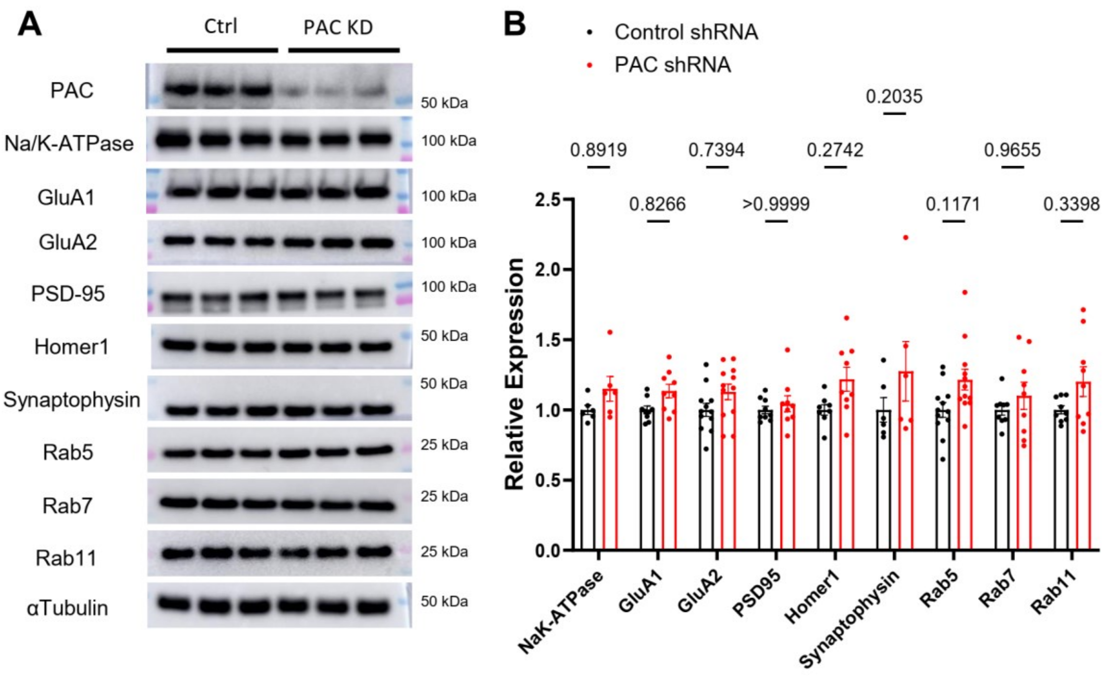


**Fig. S2. AMPARs, synaptic protein, and Rab-GTPase protein expression levels are unchanged in PAC KD neurons.**

**(A)** Representative western blot depicting protein levels of various neuronal and endosomal proteins. A polycolonal antibody targeting mouse PAC was developed and detects endogenous PAC in primary rat cortical neurons. **(B)** Quantification of protein levels normalized to alpha-Tubulin for control and PAC KD neurons.

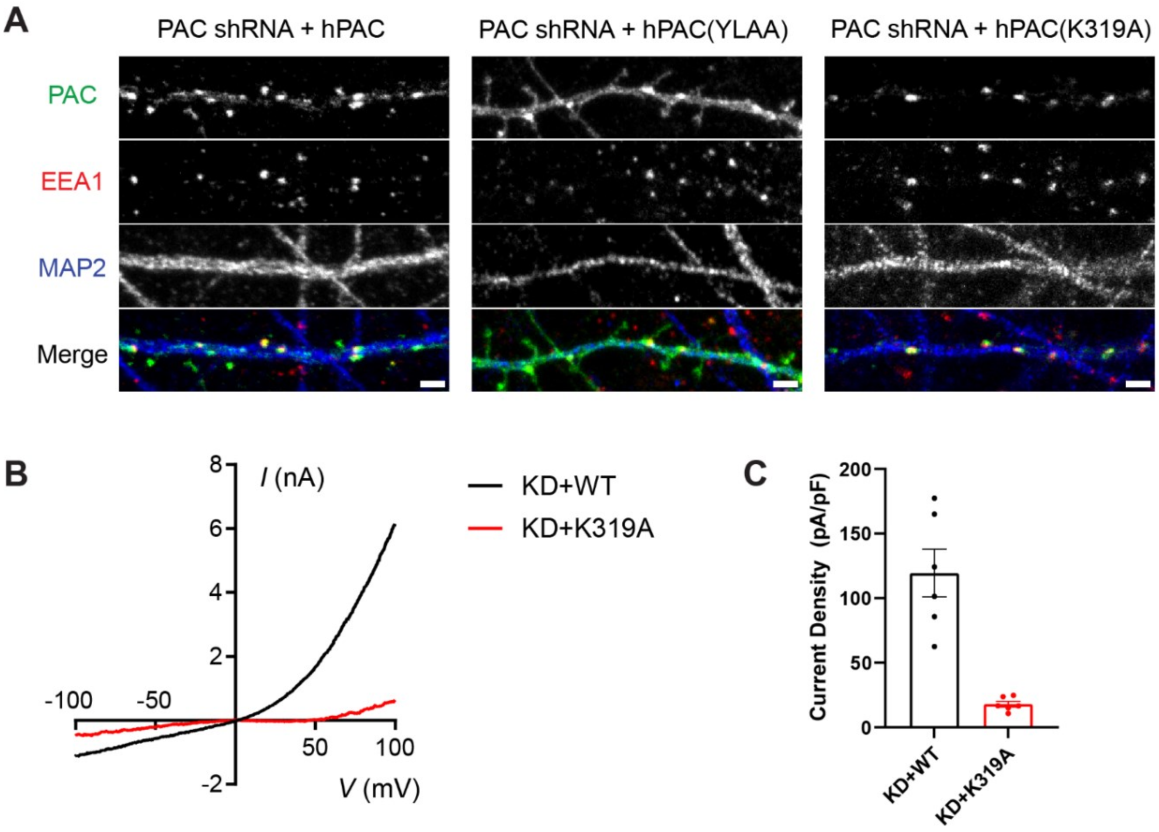


**Fig. S3. Validation of PAC mutant expression vectors.**

**(A)** Hippocampal neurons expressing vectors containing shRNA targeting rat PAC and shRNA-resistant human PAC (WT, YLAA, and K319A) immunolabeled with antibodies targeting hPAC, EEA1, and MAP2. YLAA PAC mutant displays increased surface expression compared to wild-type. PAC K319A localizes to early endosomes as expected. Scale bar 2 μm. **(B)** I-V curve of hPAC expressed in rat hippocampal neurons treated with rPAC shRNA at pH 4.6. **(C)** Quantification of current density from WT or K319A PAC at pH 4.6 in hippocampal neurons.

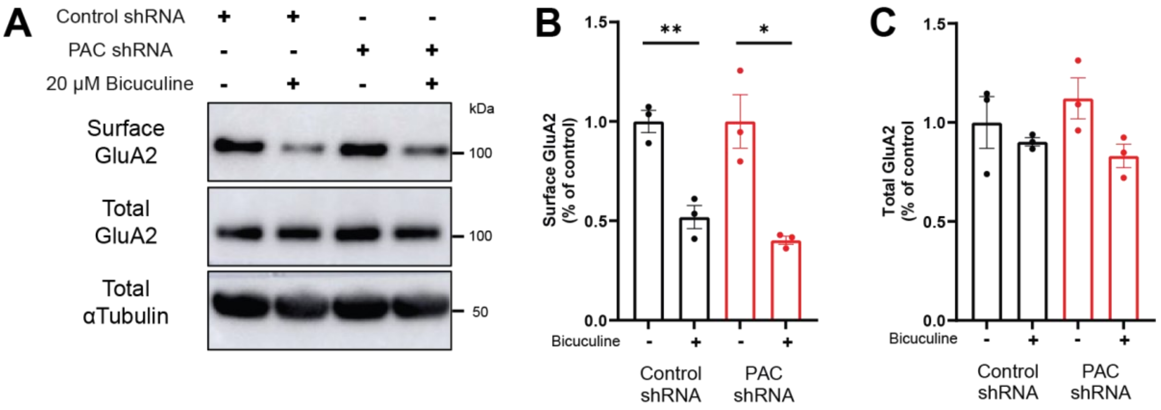


**Fig. S4. PAC deletion does not alter homeostatic downscaling.**

**(A)** Western blot depicting decrease in surface GluA2 following 48 hours of bicuculine. Both control and PAC KD neurons undergo normal homeostatic downscaling of GluA2. **(B and C)** Quantification of surface GluA2 and GluA2 protein levels after homeostatic downscaling and surface biotinylation. Error bars represent mean ± SEM. $^{**}p < 0.01$, $^{*}p < 0.05$, two-way ANOVA with Tukey correction for multiple comparisons.

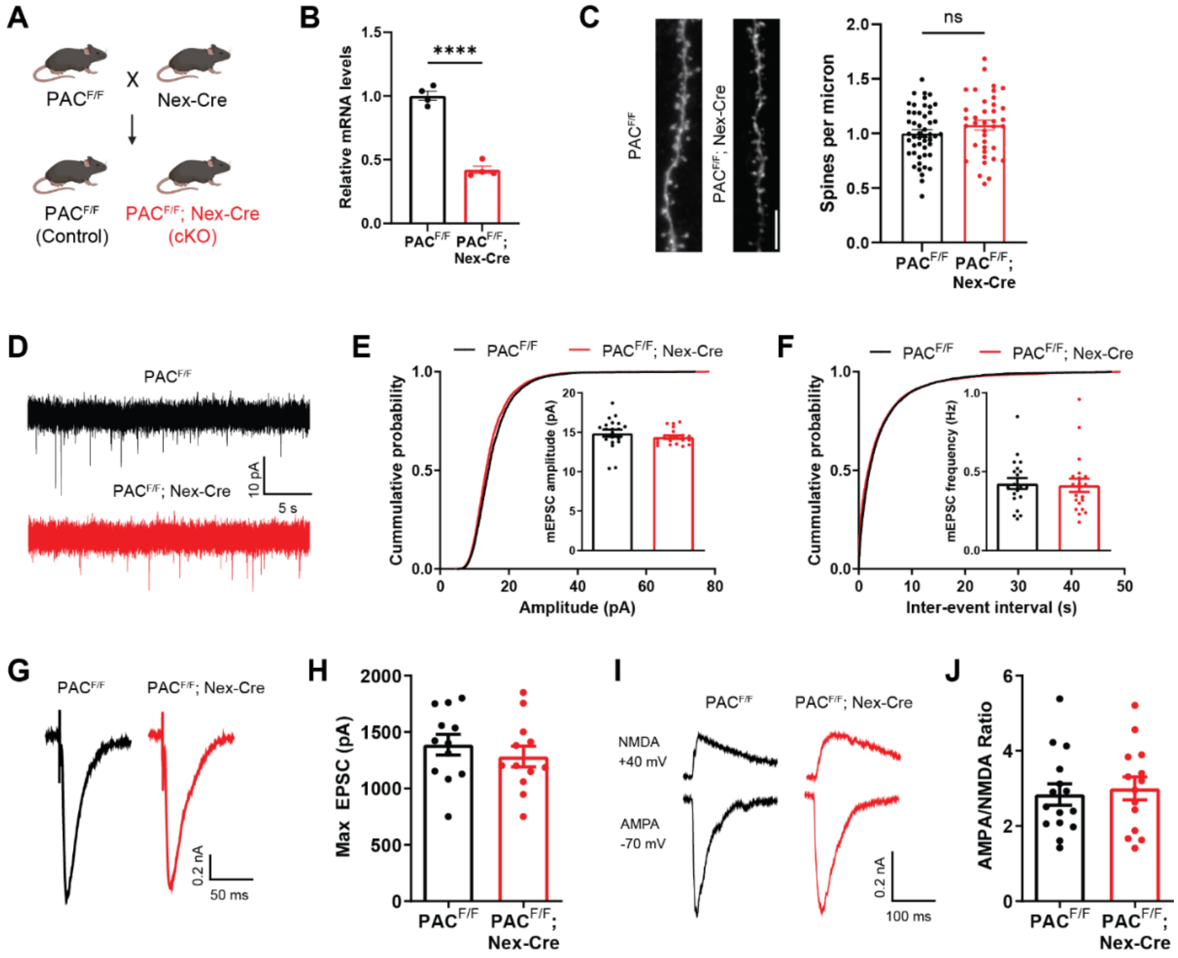


**Fig. S5. PAC cKO mice display normal synaptic transmission.**

**(A)** Breeding strategy generating neuron-specific PAC cKO mice by crossing PACF/F with the Nex-Cre line. **(B)** PAC transcript levels in whole hippocampal tissue is reduced in PAC cKO mice. **(C)** Representative images and quantification of dendritic spine density from CA1 neurons in control (n = 49 dendritic segments from 4 mice) and PAC cKO mice (n = 38 dendritic segments from 3 mice). Scale bar 5 µm. **(D)** Traces of mEPSCs in hippocampal CA1 pyramidal neurons, which are comparable between WT and cKO mice. **(E)** Quantification of frequency and cumulative probability of inter-event interval. **(F)** Quantification of mEPSC amplitude and cumulative probability of amplitude. **(G and H)** Traces and quantification of maximally evoked EPSC. **(I and J)** Traces and quantification of AMPA/NMDA ratio. Data represent mean ± SEM. ****$p<0.0001$. Data not significant unless otherwise indicated. Student's t-test for comparisons between genotypes.

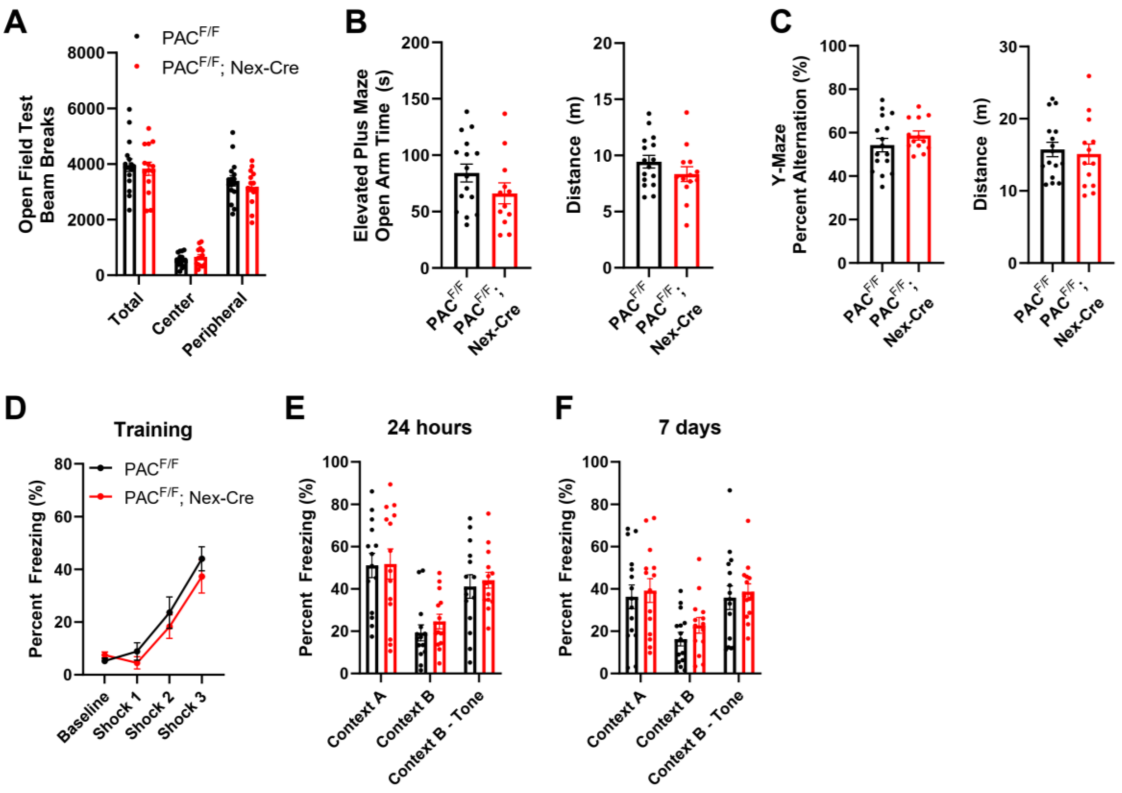


**Fig. S6. PAC cKO mice have normal locomotion, anxiety, Y-maze alternation, and fear conditioning.**

**(A)** Quantification of total, center, and peripheral beam breaks in the open field test. **(B)** Quantification of time spent in the open arm and distance traveled in the elevated plus maze. **(C)** Quantification of percent alternation and distance traveled in the Y-maze. **(D)** Quantification of percent freezing during the training period of contextual fear conditioning. **(E-F)** Quantification of percent freezing in the fear-associated context A, a novel context B, and a novel context paired with a fear-associated tone 24 hours and 7 days post training.

**Movie S1. HaloTag-GluA2 and AcidifluorORANGE allow for the visualization of AMPAR endocytosis during chemical LTD.**

Description: AMPAR internalization visualized following NMDA stimulation of primary neurons, related to Figure 4. A net increase in endocytosis does not occur in unstimulated cells, or in neurons treated with both NDMA and D-APV. PAC KD neurons do not internalize AMPARs in response to NMDA and expression of WT PAC rescues the defect of AMPAR endocytosis in a PAC KD background. YLAA and K319A PAC mutants fail to rescue AMPAR endocytosis.
